# Long identical sequences found in multiple bacterial genomes reveal frequent and widespread exchange of genetic material between distant species

**DOI:** 10.1101/2020.06.09.139501

**Authors:** Michael Sheinman, Ksenia Arkhipova, Peter F. Arndt, Bas E. Dutilh, Rutger Hermsen, Florian Massip

**Affiliations:** Theoretical Biology and Bioinformatics, Utrecht University, Padualaan 8,3584 CH, Utrecht, The Netherlands; Division of Molecular Carcinogenesis, the Netherlands Cancer Institute, Plesmanlaan 121, 1066 CX Amsterdam; Max Planck Institute for Molecular Genetics, Ihnestr. 63/73, 14195 Berlin, Germany; Berlin Institute for Medical Systems Biology, Max Delbrück Center, Berlin, Germany; Université de Lyon, Université Lyon 1, CNRS, Laboratoire de Biométrie et Biologie Evolutive UMR 5558, Villleurbanne, France

## Abstract

Horizontal transfer of genomic elements is an essential force that shapes microbial genome evolution. Horizontal Gene Transfer (HGT) occurs via various mechanisms and has been studied in detail for a variety of systems. However, a coarse-grained, global picture of HGT in the microbial world is still missing. One reason is the difficulty to process large amounts of genomic microbial data to find and characterise HGT events, especially for highly distant organisms. Here, we exploit the fact that HGT between distant species creates long identical DNA sequences in genomes of distant species, which can be found efficiently using alignment-free methods. We analysed over 90 000 bacterial genomes and thus identified over 100 000 events of HGT. We further developed a mathematical model to analyse the statistical properties of those long exact matches and thus estimate the transfer rate between any pair of taxa. Our results demonstrate that long-distance gene exchange (across phyla) is very frequent, as more than 8% of the bacterial genomes analysed have been involved in at least one such event. Finally, we confirm that the function of the transferred sequences strongly impact the transfer rate, as we observe a 3.5 order of magnitude variation between the most and the least transferred categories. Overall, we provide a unique view of horizontal transfer across the bacterial tree of life, illuminating a fundamental process driving bacterial evolution.

## 1. Introduction

Microbial genomes are subject to loss and gain of genetic material from other organisms [5, 60], via a variety of mechanisms: conjugation, transduction, and transformation, collectively known as horizontal gene transfer (HGT) [70, 26]. The exchange of genetic material is a key driver of microbial evolution that allows rapid adaptation to local niches [6]. Gene acquisition via HGT can provide microbes with adaptive traits, conferring a selective advantage in particular conditions [34, 43], and eliminates deleterious mutations, resolving the paradox of Muller’s ratchet [71].

Since the discovery of HGT more than 50 years ago [24] many cases of HGT have been intensively studied. Several methods to infer HGT rely on identifying shifts in (oligo-)nucleotide compositions along genomes [63]. Other methods are based on discrepancies between gene and species distances, *i*.*e*., surprising similarity between genomic regions belonging to distant organisms that cannot be satisfactorily explained by their conservation [38, 50, 35, 52, 18, 19, 9]. For example, genomes from different genera are typically up to 60 – 70% identical, meaning that one in every three base pairs is expected to differ. The presence of regions in different genomes that are significantly more similar than expected can be interpreted as evidence of recent HGT events. Using such methods the transfer of drug- and metal-resistance genes [31], toxin-antitoxin systems [73] and virulence factors [22, 51] have been observed numerous times. It is also known that some bacterial taxa, such as members of the family of *Enterobacteriaceae* [20], are frequently involved in HGT, whereas other groups, such as extracellular pathogens from the *Mycobacterium* genus [21], rarely are. Notably, the methods used in the detection and analysis of instances of HGT are computationally complex and can be used to discover HGT event in at most hundreds of genomes simultaneously. Consequently, a general overview of the diversity and abundance of transferred functions, as well as the extent of involvement across all known bacterial taxa in HGT, is still lacking. In particular, exchanges of genetic material between distant species – because discovering such long-distance transfers requires the application of computationally costly methods to very large numbers of genomes – are rarely studied.

In this study we use a novel approach to address these questions. Our method is based on the analysis of long exact sequence matches found in the genomes of distant bacteria. Exact matches can be identified very efficiently using alignment-free algorithms [17], which makes the method much faster than previous methods that rely on alignment tools. This allows us to study transfer events between 1 343 042 bacterial contigs, belonging to 93 481 genomes, encompassing a total of 0.4 Tbp. We identified all long exact matches shared between bacterial genomes from different genera. Such long matches are unlikely to be vertically inherited, and we therefore assume that they result from HGT.

In a quarter of all bacterial genomes, we detected HGT across family borders, and 8% participated in HGT across phyla. This shows that genetic material frequently crosses distant taxonomic borders. The length distribution of exact matches can be accounted for by a simple model that assumes that exact matches are continuously produced by transfer of genetic material and subsequently degraded by mutation. Fitting this model to empirical data allow us to estimate the effective rate at which HGT generates long sequence matches in distant organisms. Furthermore, the large number of transfer events identified allows us to conduct a functional analysis of horizontally transferred genes.

## 2. Results

### 2.1. HGT detection using exact sequence matches

We identified HGT events between distant bacterial taxa by detecting long exact sequence matches shared by pairs of genomes. We exploit that, in phylogenetically distant genome pairs, sequences that are shared by both genomes due to linear descent (orthologous sequences) have low sequence identity. Therefore long sequence matches in such orthologs are exceedingly rare. Generally, the average nucleotide sequence identity between bacterial genomes selected from different genera is at most 60 to 70% [61]. In the absence of HGT, the probability of observing an exact match longer than 300 bp between a given pair of genomes is then extremely small (of the order of 10^−40^ if we assume uniform divergence along the genomes). Thus, even if millions of genome pairs with such divergence are analysed, the probability to observe even one such a match remains extremely low, such that one does not expect to find a single hit of this size between any two bacterial genomes by chance.

Fig. 1 illustrates this point. In the dot-plot comparing the genome sequences of two *Enterobacteriaceae, Escherichia coli* and *Salmonella enterica* (Fig 1A), we observe numerous exact matches smaller than 300bp along the diagonal, revealing a conservation of the genomic architecture at the family level. Filtering out matches shorter than 300bp (Fig 1B) completely removes the diagonal line, confirming that exact matches in the orthologous sequences of these genomes are invariably short.

**Figure 1:**
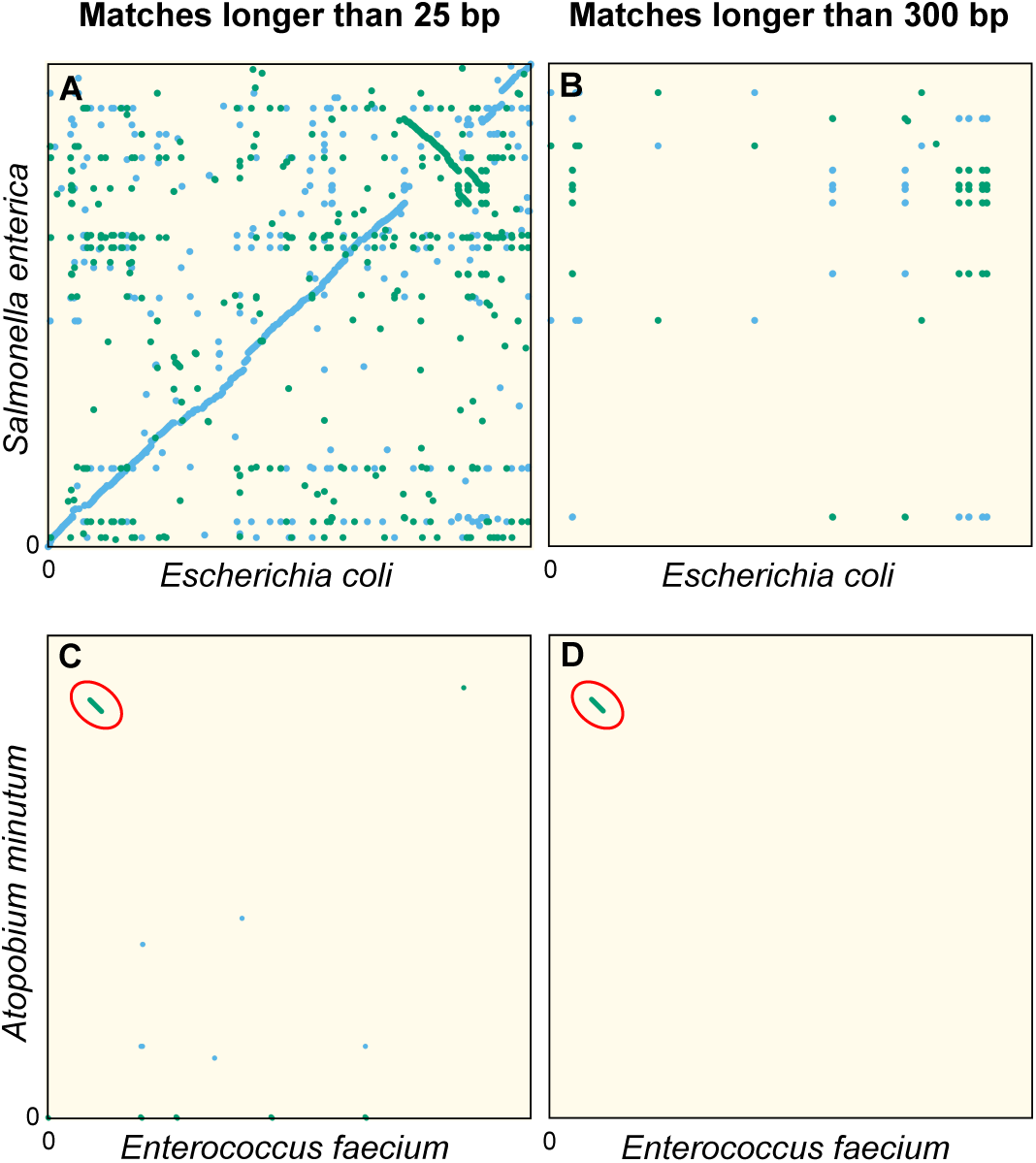
Dot plots of the exact sequence matches found in different pairs of distant bacteria. Matches are indicated as lines drawn with a wide stroke. Thus, short matches appear as dots. Blue lines indicate matches in the forward strand, green lines those in the reverse complement. **(A-B)** Full genomes of *Escherichia coli K-12 substr. MG1655* (U00096.3) and *Salmonella enterica* (NC_003198.1), which both belong to the family of *Enterobacteriaceae*. Panel A shows all matches longer than 25 bp. The sequence similarity and synteny of both genomes, by descent, is evident from the diagonal blue line. Panel B only shows matches longer than 300 bp. **(C-D)** Same as Panels A-B, but for the first 1.4 Mbp of *Enterococcus faecium* (NZ_CP013009.1) and *Atopobium minutum* (NZ_KB822533.1), which belong to different phyla, showing few matches longer than 25 bp (Panel C). Yet, a single match of 19 117 bp is found, as indicated with red ellipses in Panels C-D. The most parsimonious explanation for this long match is an event of horizontal gene transfer.

Because very long exact sequence matches are extremely unlikely in orthologs, those that do occur are most likely xenologs: sequences that are shared due to relatively recent events of HGT. As an example, Fig. 1C shows a dot plot comparable to Panel 1A, but now comparing the genomes of *Enterococcus faecium* and *Atopobium minulum*. No diagonal line is seen because these genomes belong to different phyla and therefore have low sequence identity. Nevertheless, an exact match spanning 19 117 bp is found (diagonal green line highlighted by a red ellipse). The most parsimonious explanation for such a long match is a recent HGT event. In addition, the GC content of the match (55%), deviates strongly from that of both contigs (38.3% and 48.9%, respectively), another indication that this sequence originates from an HGT event [63]. Comparing the sequence of this exact match with all non-redundant GenBank CDS translations using `blastx` [1] we find very strong hits to VanB-type vancomycin resistance histidine, antirestriction protein (ArdA endonuclease), and an LtrC-family phage protein that is found in a large group of phages that infect Gram-positive bacteria [62]. Together, this suggests that the sequence was transferred by transduction and established in both bacteria aided by natural selection acting on the conferred vancomycin resistance.

In the following we assume that long identical DNA segments found in pairs of bacteria belonging to different genera reveal HGT. We stress, however, that a matching sequence may not have been transferred directly between the pair of lineages in which it was identified: more likely, it arrived in one or both lineages independently, for instance carried by a phage or another mobile genetic element that transferred the same genetic material to multiple lineages through independent interactions.

In the following, we restrict our study to matches longer than 300 bp to minimise the chance that those matches result from vertical inheritance. Because transferred sequence accumulate mutations, matches longer than 300 bp must originate from relatively recent events. Assuming a generation time of 10 hours [28], we estimate the detection horizon to be of the order of 1000 years ago (see Methods).

### 2.2. Empirical length distributions of exact matches obey a power law

To study HGT events found in pairs of genomes from different genera, we considered the statistical properties of *r*, the length of exact matches. To do so, we selected all bacterial genome fragments longer than 10^5^ bp from the NCBI RefSeq database (1 343 042 in total), and identified all sequence matches in all pairs of sequences belonging to different genera (≈ 10^9^ pairs). We then analysed the distribution of the match lengths found, called the match-length distribution or MLD. A comparable approach has previously been applied successfully to analyse the evolution of eukaryotic genomes [25, 44, 45, 46].

While the vast majority of matches is very short (< 25 bp), matches with a length of at least 300 bp do occur and contribute a thick tail to the MLD (Fig. 2). Strikingly, over many decades this tail is well described by a power law with exponent −3:

**Figure 2:**
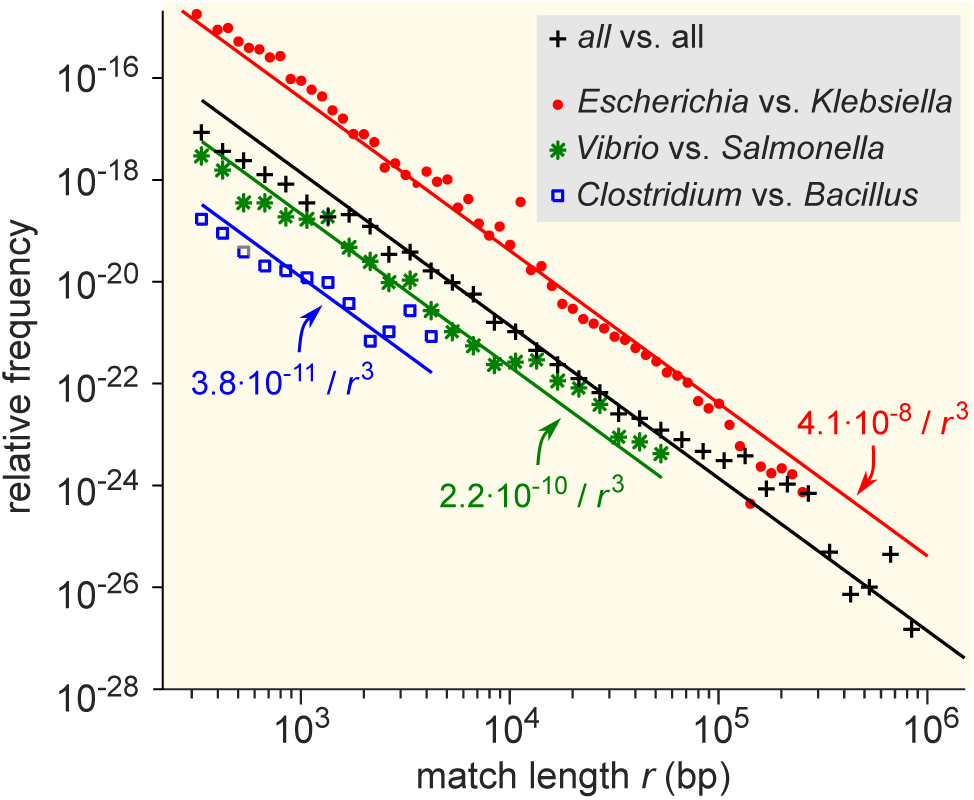
Match length distributions (MLDs) obtained by identifying exact sequence matches in pairs of genomes from different genera, based on matches between *Escherichia* and *Klebsiella* (red dots), *Vibrio* and *Salmonella* (green stars), and *Clostridium* and *Bacillus* (blue squares). Black plus signs represent the MLD obtained by combining the MLDs for *all* pairs of genera. Each MLD is normalised to account for differences in the number of available genomes in each genus (see Methods). Only the tails of the distributions (length *r* ≥ 300) are shown. Solid lines are fits of power-laws with exponent –3 (Eq. (1)) with just a single free parameter.

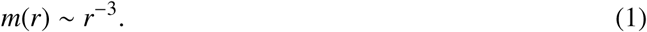

The same power law was found if the analysis was restricted to matches between genomes from two particular genera (Fig. 2).

Note that the number of long matches found in a single pair of genomes is usually very small, prohibiting a statistical analysis of their match length distribution. Hence, in this study we conduct all statistical analyses at the level of genera. The MLD for a pair of genera *G*_1_ and *G*_2_ is defined as the normalized length distribution of the matches found in all pairwise comparisons of a contig from *G*_1_ and a contig from *G*_2_ (see Methods).

### 2.3. A simple model of HGT explains the power-law distribution of exact sequence matches

A simple model based on a minimal set of assumptions can account for the observed power law in the MLD. Let us assume that, due to HGT, a given pair of bacterial genera A and B obtains new long exact matches at a rate ρ, and that these new matches have a typical length *K* much larger than 1 bp. These matches are established in certain fractions *f*_A_ and *f*_B_ of the populations of the genera, possibly aided by natural selection. Subsequently, each match is continuously broken into shorter ones due to random mutations that happen at a rate μ per base pair in each genome. Then the length distribution of the broken, shorter matches, resulting from all past HGT events, converges to a steady state that for 1 bp ≪ *r* < *K* is given by the power law *m*(*r*) = *A*/*r*^−3^, with prefactor:

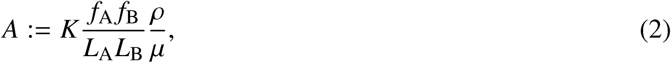

consistent with Eq. (1). Here *L*_A_ (resp. *L*_B_) is the average genome length of all species in genus A (resp. B); see Methods. Hence, the power law observed when analysing pairs of bacterial taxa can be explained as the combined effect of many HGT events that occurred at different times in the past. While the model above makes several strongly simplifying assumptions, many of these can be relaxed without affecting the power-law behaviour; see Methods for an extended discussion.

In the model, the prefactor *A* quantifies the abundance of long exact matches and hence is a measure of the rate with which two taxa exchange genetic material. Eq.2 shows that *A* reflects the bare rate of the transfer events, the typical length of the transferred sequences, as well as the extent to which the transferred sequences are established in the receiving population, possibly aided by selection. By contrast, because of the normalisation of the MLD (see Methods), *A* does not scale with the number of genomes in the genera being compared and is thus robust to sampling noise, so that the value of *A* can be used to study the variation in HGT rate between genera.

### 2.4. Long-distance gene exchange is a widespread mechanism in the bacterial domain

The analysis above has allowed us to identify a large number of HGT events. In addition, the derivations in the previous section provide a method to quantify the effective HGT rate between any two taxa by measuring the prefactor *A*. As Supp. file 1 (resp. Supp. file 2), we provide the value of *A* for all pairs of families (genera). Using these methods, we then studied the HGT rate between all pairs of bacterial families in detail.

Fig. 3 plots the prefactors *A* for all pair of families. Families for which the available sequence data totals less than 10^7^ bp were filtered out since in such scarce datasets, typically no HGT is detected (Fig. S1), and the prefactor cannot reliably be estimated (see Supp. File 3 for the total length of all families). A first visual inspection of the heatmap reveals that the HGT rate varies drastically (from 10^−^16 to 10^−^8) from one pair to another (Fig. 3). First, the large squares on the diagonal of the heatmap indicate that HGT occurs more frequently between taxonomically related families. This is especially apparent for well-represented phyla including *Bacteriodetes, Proteobacteria, Firmicutes*, and *Actinobacteria*. Yet, we also observe a high transfer rate between many families belonging to very distant phyla, indicating that transfer events across phyla are also frequent. Notably, we find that some families present a highly elevated HGT rate across the phylogeny; these families are visible in the heatmap (Fig. 3) as long bright lines, both vertical and horizontal.

**Figure 3:**
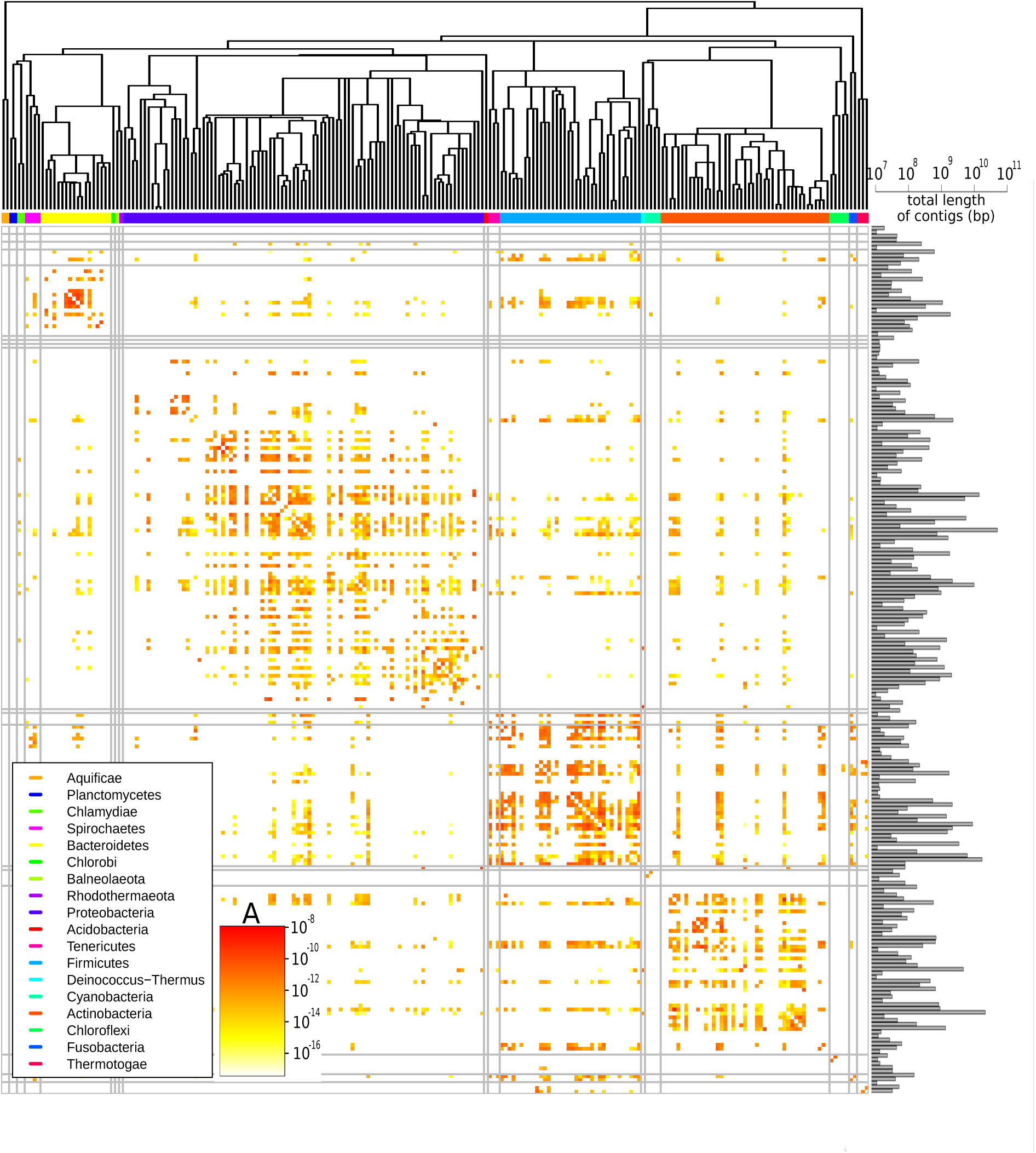
Effective pairwise HGT rate at the family level. For each pair of families the prefactor *A* is displayed (decimal logarithmic scale, see colorbar and Supp. file 1). The phylogenetic tree of bacterial families, taken from [37], is shown at the top. Phyla are indicated with coloured bars next to the upper axes of the heatmap (see legend). On the diagonal the values are set to zero. Black vertical and horizontal lines represent borders between phyla. The barplot on the right side of the heatmap shows the cumulative genome sizes of each family (decimal logarithmic scale).

We studied the HGT rate variations in more detail in a restricted dataset which included only long contigs (> 10^6^ bp) to reduce the risk of potential artefacts (see Methods). This dataset still comprises 138, 273 matches longer than 300bp.

The analysis of the restricted dataset reveals the extent of HGT in bacteria, even between distant species (Fig. 4). Indeed, we find that 32.6% of species have exchanged genetic material with a species from a different family in the last ∼ 1000 years. Moreover, we find that 8% of species have exchanged genetic material with a species from a different phylum. Finally, the species involved in these distant exchanges are spread across the phylogenetic tree: the species involved in long-distance transfers belong to 19 different phyla (out of 34).

**Figure 4:**
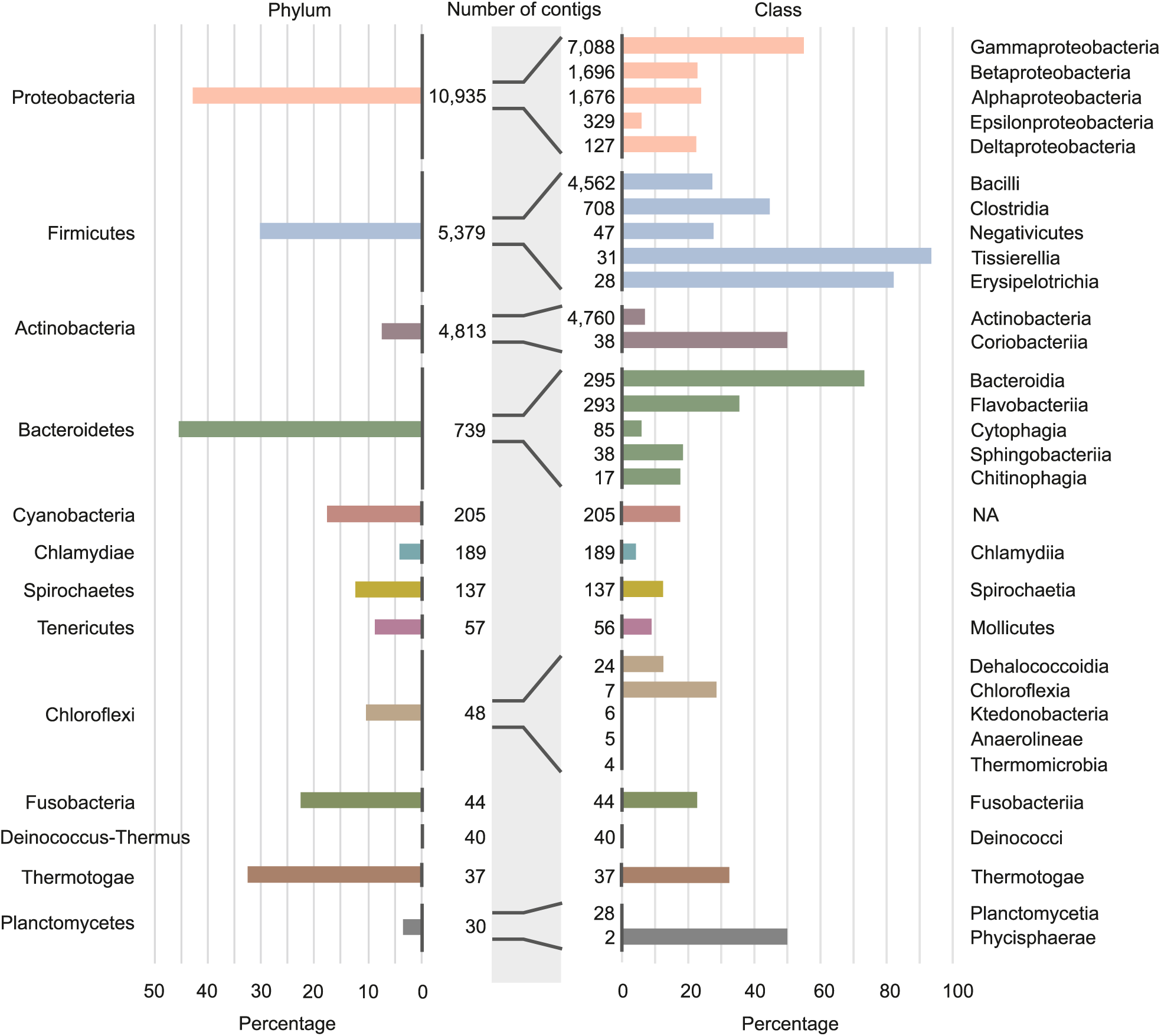
Involvement of different phyla and classes of bacteria long-distance HGT. Percentage of contigs involved in at least one long distance HGT event grouped at phylum level (left panel) and at classes level (right panel). Note that only the classes with the largest numbers of contigs are shown in the figure (see Supp. file 4 for all data). Numbers of contigs belonging to the phyla and classes are given in the middle part of figure.

The data also unveil that the propensity of species to exchange genetic material is very heterogeneous, and varies dramatically between closely related classes. For instance, within the phylum *Firmicutes*, we find classes in which we detected HGT in only a small percentage of species (30% in the *Negativicutes*), while in other classes we find events in almost all species (> 90% in *Tissierellia*, Fig. 4 and Supp. file 4). This trend can be observed in most of the phyla and raises the question of which species features drive HGT rate variations.

### 2.5. The rate of HGT decreases with taxonomic distance

To better understand the causes of the large variations in transfer rate between different families, we next studied the effect of biological and environmental properties on the HGT rate.

First, we assessed the impact of the taxonomic distance between genera on the HGT rate. To do so, we computed the prefactor *A* for pairs of genera at various taxonomical distances (Fig 5). On average this prefactor decreases by orders of magnitude as the taxonomic distance between the genera increases (inset of Fig 5). In particular, the average prefactor obtained when considering genera from the same family is more than three orders of magnitude higher than when considering genera from different phyla. These results support the notion that the divergence between organisms plays an important role in the rate of HGT between them [53, 7, 49, 27, 12, 15, 2] (see also Fig. S2). Note however that a lower effective rate of HGT can be due to a lower transfer rate of genetic material and/or a more limited fixation in the receiving genome, and the model cannot distinguish those two scenarios.

**Figure 5:**
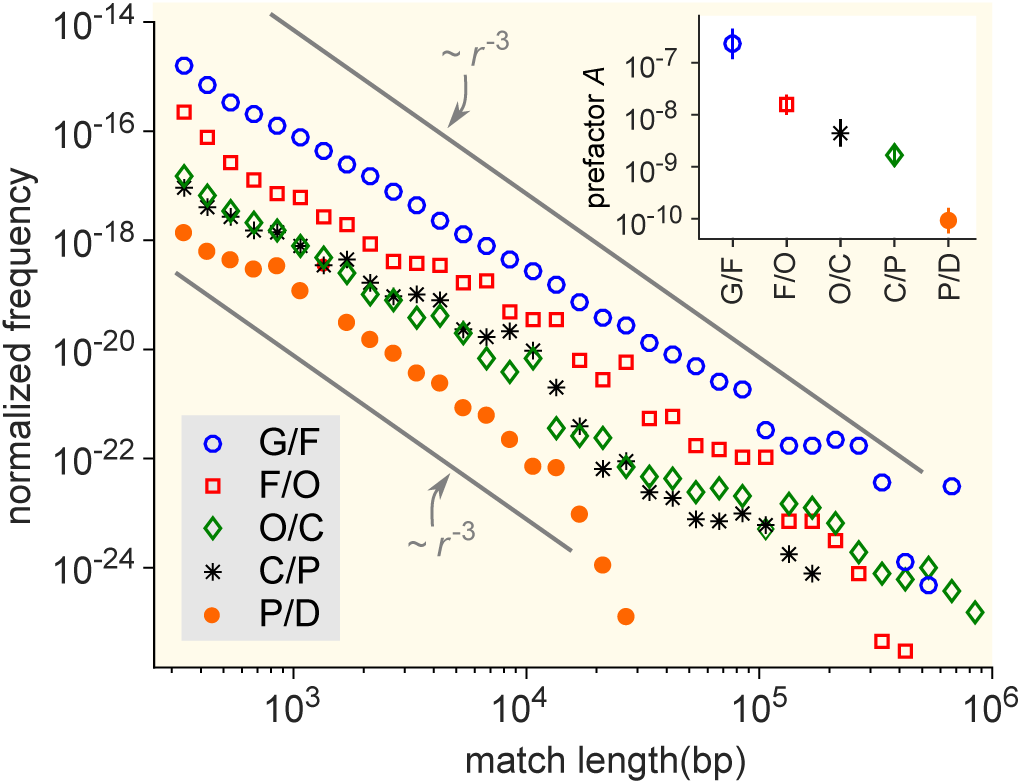
Distribution of matches lengths resulting from comparison of genera at a given taxonomic distance. G/F (blue circles): All pairwise comparison of genera from the same family; F/O (red squares) matches between genera of different families but in the same order; O/C (green diamonds), different orders but same class; C/P (black stars) different classes but same phylum; P/D (red circles) different phylum but same domain. The gray lines indicate the power-law dependence *m*(*r*) = *Ar*^−3^. Inset: prefactor *A* for each of the distributions in the main figure. The prefactor decrease by orders of magnitude as the taxonomic distance increases.

To further explore the factors that influence the value of *A* we calculated MLDs for sets of genera from different ecological environments: gut, soil, or marine (Fig. S3), regardless of their taxonomic distance. Our results suggest that the effective rate of HGT is about 1 000 times higher among gut bacteria than among marine bacteria. This pattern is observed both for the rates of HGT within ecological environments (*i.e*., HGT among gut bacteria vs. among marine bacteria) and the rates of crossing ecological environments (*i.e*., HGT between gut and soil bacteria versus between marine and soil bacteria). The soil bacteria take an intermediate position between the gut and the marine bacteria. Moreover, bacteria from the same environment tend to share more matches than bacteria from different environments, consistent with previous analyses [69].

A similar analysis demonstrates that the HGT propensity among gram-positive bacteria and among gram-negative bacteria is much larger than between these groups (see Fig. S4). The groups of bacteria with GC poor and GC rich genomes exhibit a similar pattern (see Fig. S5). We note however, that all these factors correlate with each other [30]. From our analysis, the contribution of each factor to the effective rate of HGT therefore remains unclear.

### 2.6. Large variation in the HGT rates between different categories of genes

To better understand the factors that explain variations in observed HGT rates, we next conducted a functional analysis of transferred sequences. To functionally annotate the transferred sequences, we first queried twelve databases, each specifically dedicated to genes associated with a particular function (see Table S1). Comparing to a randomised set of sequences (see Methods) reveals that the gene functions of the transferred sequences strongly impact the transfer rate, as we observe a 3.5 order of magnitude variation between the most and the least transferred categories (Fig. 6 and Table S1).

**Figure 6:**
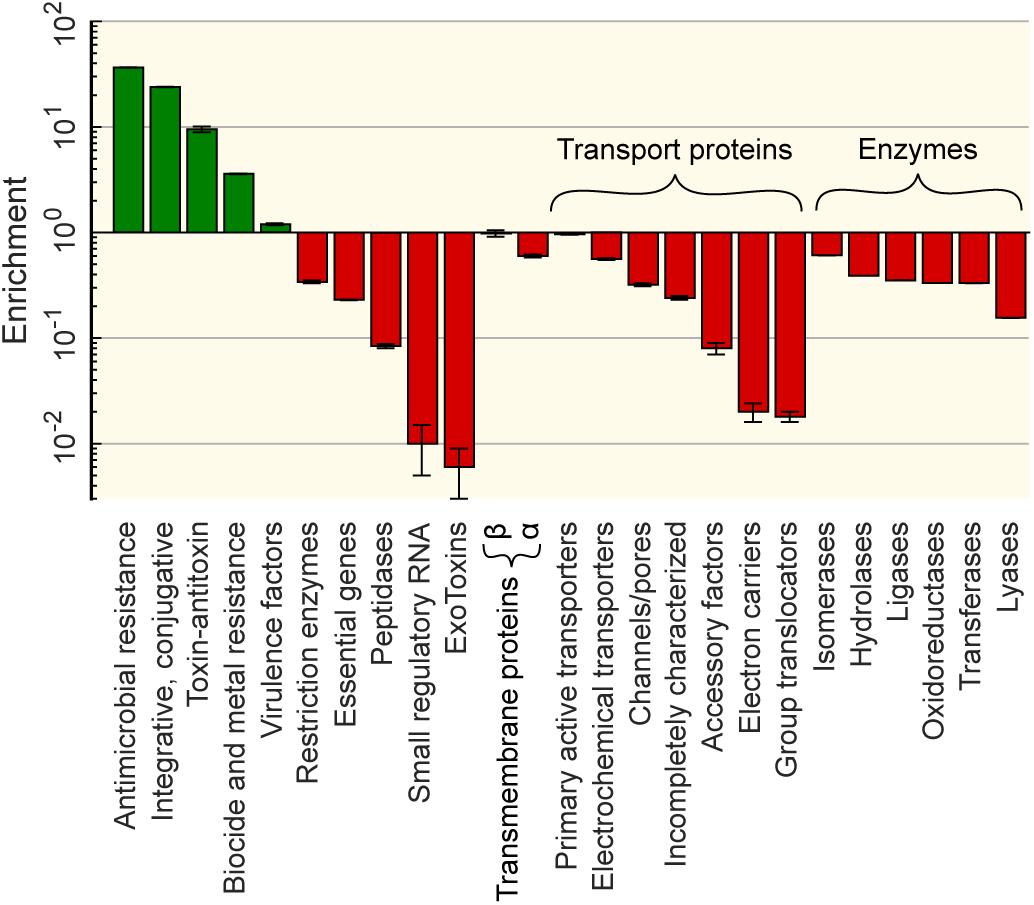
Functional enrichment of the sequences involved in HGT. Enrichment for each gene category (vertical axis) are computed relative to the random control for each set of genes from a certain category from the appropriate database (see Methods). Enrichment for gene resistance against different types of antibiotics and different biocides can be found in Supp. file 5.

More specifically, antibiotic and metal resistance genes are among the most widely transferred classes of genes (resp. 37× and 4× enrichment compared to random expectation), in good agreement with previous evidence [31, 74, 23]. The enrichment of resistance genes is expected since their functions are strongly beneficial for bacterial populations under specific, transient conditions. Interestingly, genes providing resistance against tetracycline and sulfonamide antibiotics — the oldest groups of antibiotics in use — are the most enriched (see the full list in Supp. file 5). In addition, we also find a strong enrichment among the transferred genes of genes classified as integrative and conjugative elements, suggesting that these genes mediated the HGT events [57, 49]. In contrast, exotoxins and small regulatory RNAs are the least transferred genes (≈ 100× depletion). More generally, genes in the wider “Transport proteins” and “Enzymes” categories are strongly underrepresented in the detected HGT events.

To obtain a better understanding of the function of the transferred sequences, we also annotated the transferred sequences using SEED Subsystems [55] (Methods). While the 12 curated databases queried above are more complete and accurate on their specific domains, using the SEED Subsystem allows to test for over- or underrepresentation of a broader set of functions. The results of this second method are in good agreement with the database queries as the broad categories linked to “Phages, Prophages, Transposable elements, Plasmids”, and to “Virulence, Disease and Defense” are found to be the most enriched, although with a smaller enrichment (4.3 and 2.5 fold enrichment respectively, see Supp. file 6).

In addition to previously known enriched functions, we also discovered a strong enrichment (2.8×compared to the control, conditional test adjusted *p*-value < 10^−16^, see Methods) for genes in the “iron metabolism” class. Indeed, a wide range of iron transporters, parts of siderophore and enzymes of its biosynthesis appeared in our HGT database, in line with previous analysis focusing on cheese microbial communities [4]. Hence, the results show that the horizontal transfer of genes related to iron metabolism occurs in a wide set of species and is not restricted to species found in cheese microbial communities. Notably, the proteins in the “iron metabolism” functional category can be identified in transferred sequences belonging to 6 different bacterial phyla.

Among the enriched SEED subsystem categories, another interesting example is the enrichment for genes in the “flagellar motility” category (5.49× compared to the control, conditional test adjusted *p*-value < 10^−16^). The flagellum is a complex multi-protein locomotor organ of bacteria [42] that has been found even in non-motile species [29]. An interesting feature of flagella is their own protein export system, which enables transfer of extracellular flagellar proteins outside of bacterial cells [48]. We found that the exact matches code for a set of proteins of this export system. This result is supported by the recent finding that flagellin glycosilation islands can be transferred [16]. The frequent transfer of flagellar genes, combined with the fact that flagellar genes have been found even in non-motile species [29] could indicate that the export system of the flagella takes part in transport of other compounds, such as toxic chemicals.

Overall, the above findings confirm the strong enrichment of resistance genes among HGT events and validate the good resolution of our methods, and its power at shedding a new light on the properties of horizontal gene transfer.

## 3. Discussion

In this study, we developed a computationally efficient method to identify recent HGT events. This method provided an unprecedentedly large database of horizontal gene transfer events between any two genera in our database of 93 481 organisms. Our analysis reveals that HGT between distant species is extremely common in the bacterial world, with 32.6% percentage of organisms having taken part in an event that crossed genus boundaries in the last ∼ 1000 years. While a similar analysis has been conducted on a much smaller dataset (about 2, 300 organisms) Smillie et al. [69], this study is, to our knowledge, the first to provide an extensive description of HGT in the microbial world at this scale.

One striking result of our analysis is the finding that HGT is also common between very distant organisms. Indeed, 8% of the organisms we studied have been involved in a transfer of genetic material with at least one bacteria from another phylum in the last ∼ 1000 years. The molecular mechanisms at play in these long distance transfer events remain to be elucidated, for instance via a dedicated study targeting families with very high exchange rate we identified. Analysing the statistical properties of the exact sequence matches in distantly related genera, we were able to quantify the effective rate of HGT for different comparisons (See Supp. File 1 and 2 for an estimation of all pairwise HGT rate at the family and at the genera level). Doing so, we find that the HGT rate varies dramatically between families (Fig. 3), posing the question of the factors influencing the HGT rate. Our study confirms that the HGT rate decreases with the divergence between the two bacteria exchanging material (Fig. 5 and Supp. Fig. S2), and is larger for pairs of bacteria with similar properties, such as ecological environment, GC content and Gram staining (Fig. S3, S4 and S5). However, since all these properties are correlated with each others, we could not disentangle the independent contribution of each of those features to the HGT rate.

Finally, our functional analysis of the transferred sequences shows that the function of a gene also strongly influences its chance of being exchanged (Fig. 4). As expected, genes conferring antibiotic resistance are the most widely transferred. In contrast, some functional categories are strongly underrepresented in the pool of transferred genes. For instance, genes that are involved in transcription, translation, and related processes as well as those involved in metabolism are all depleted in our HGT database. One potential explanation could be that these genes generally co-evolve with their binding partners [33]. As such, their transfer would be beneficial to the host species only if both the effector and its binding partner were to be transferred together. As simultaneous HGT of several genes from different genome loci is very unlikely (unless they are co-localized), these genes are not prone to HGT. In addition, transcription, translation, and related processes are core functions that are ubiquitous in bacterial species. As such, transfer of such genes is unlikely to grant the receiving species a new function. Hence, transfers of housekeeping gene are less likely to confer a strong evolutionary advantage, which could explain their under-representation in the HGT dataset.

We found that the tail of the MLD follows a power law with exponent −3. This observation is particularly robust both empirically and theoretically. Indeed, the empirical MLDs we observe span between 2 and 4 orders of magnitude, with an exponent always equal or close to –3. In addition, many of the simplifying assumptions of the model can be relaxed without breaking the specific power-law behaviour, provided that HGT events have taken place continuously and at a non-zero rate up to the present time (see Methods). Whether HGT is a continuous process on evolutionary time scales or instead occurs in bursts has been a matter of debate [65, 33, 76], and burst of transfer event at some point in the past might explain some of the deviations from the –3 power-law behaviour we observe (Fig. 5). In addition to HGT bursts, other complex evolutionary mechanisms that we do not consider in our model could in theory explain those deviations, including mechanisms of gene loss that allow bacteria to eliminate detrimental genes, or selfish genetic elements [72]. Finally, misclassifications of contigs as well as errors in genome assembly could bias the estimation of the effective HGT rate *A*.

Although it is widely accepted that bacteria often exchange their genes with closely related species [2] via HGT, our large-scale analysis of HGT shed new light on gene exchange in bacteria. Our analysis indicates that near 8% of all sequenced bacterial genomes share genes with at least one bacteria from other phylum, revealing the true scale of long distance gene transfer events. Evidently, long-distance exchange of genetic material is a recurrent and wide spread process, with specific statistical properties, suggesting that horizontal gene transfer plays a decisive role in maintaining the available genetic material throughout evolution.

## 4. Methods

### 4.1. Identification of exact matches

Reference bacterial sequences [54] were downloaded from the NCBI FTP server on 3 April 2017 together with taxonomy tree files. We identified maximal exact matches using the MUMmer [17] software with the maxmatch option, which finds all the matches regardless of their uniqueness.

### 4.2. Empirical calculation of the MLD for pairs of genera and sets of genera

To analyse MLDs, we use all contigs longer than 10^5^bp. The MLD of a pair of genera *i* and *j* is defined as

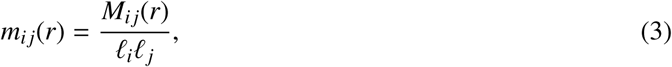

where *M*_*i j*_(*r*) is the number of matches of length *r* between all contigs of genus *i* and all contigs of genus *j*. 𝓁_*x*_ is the total length of the available contigs of genus *x*. The expected number of matches found in the analysis of a pair of genera scales with the amount of sequence data available for these genera. Normalising by 𝓁_*i*_ *𝓁*_*j*_ ensures that *m*_*i j*_(*r*) does not scale with the database size, so that the *m*_*i j*_(*r*) for different pairs of genera can be compared.

In Fig. 2, 5 and S2, S3, S4, S5 we show MLDs based on the matches found between pairs of sequences from two sets of genera. These MLDs were calculated as follows:

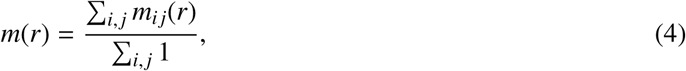

where the index *i* runs over the genera from the first set and the index *j* runs over the genera from the second set.

### 4.3. Analytical calculation of the MLD predicted by a simple model of HGT

A simple model based on a minimal set of assumptions can account for the observed power-law distributions. We first consider a particular event of HGT in which two bacterial genera gain a long exact match of length *K* ≫ 1 via HGT. After time *t*, the match is established in certain fractions of the populations of both genera, denoted *f*_1_ and *f*_2_, respectively, possibly aided by natural selection. By this time, the match is expected to be broken into shorter ones due to random mutations, which we assume occur at a constant effective rate *μ* = (*μ*_1_ + *μ*_2_)/2 at each base pair, where μ_1_ and μ_2_ are the mutation rates of genus 1 and 2.

Suppose that we now sample *n*_1_ genomes from genus 1 and *n*_2_ from genus 2 and calculate the MLD according to equation 3. Then in the regime 1 ≪ *r* < *K* the contribution of the matches derived from this particular HGT event is given by [78, 44]:

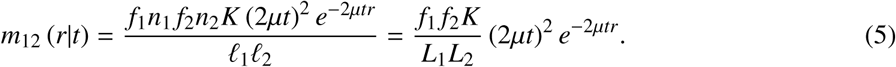

Here, *L*_1_ and *L*_2_ are the average lengths of the genomes sampled from the two genera. Equation 5 shows that each individual HGT event contributes an exponential distribution to the MLD.

The full MLD is composed of contributions of many HGT events that happened at different times in the past. Assuming a constant HGT rate ρ, the HGT events are uniformly distributed over time, which results in the following full MLD [45]:

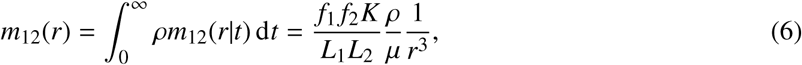

which yields the observed power-law with exponent −3.

The prefactor

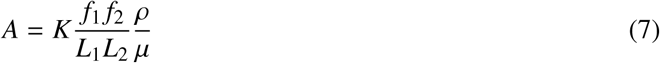

in Eq. (1) can be interpreted as an effective transfer rate per genome length. It depends on several parameters: the transfer rate from one species to another per genome length *ρ*/(*L*_1_*L*_2_), the length of the transferred sequences *K*, the degree to which the sequence is establishment in the population of the two genera *f*_1_ and *f*_2_, and the effective mutation rate *μ*.

To fit the power law (1) to the empirical data, we binned the tail (*r* > 300) of the empirical MLD (using logarithmic binning), and then applied a linear regression with a fixed regression slope of −3 and a single fitting parameter, *i.e*., the intercept ln(*A*).

### 4.4. Robustness of the power-law behaviour

For simplicity, the above argument makes several strong assumptions, including that *μ, K, f*_1_ and *f*_2_ are the same for all HGT events and that these events are distributed uniformly over time. However, if these assumptions are relaxed the power law proves to be remarkably robust. First, we could assume that all of the above parameters differ between HGT events, according to some joint probability distribution *P*(*K, μ, f*_1_, *f*_2_). As long as this distribution itself does not depend on the time *t* of the event, equation 6 then becomes

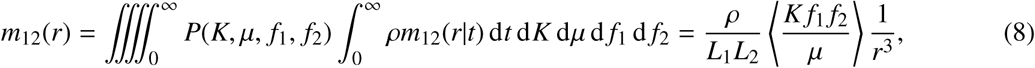

where the angular brackets denote the expectation value. The power law remains, except that the prefactor now represents an average over all possible parameter values. Second, we can relax the assumption that the divergence time *t* is uniformly distributed (*i.e*., that HGT events were equally likely at any time in the past).

In general, equation 6 should then be replaced by

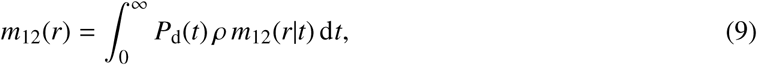

in which *P*_d_(*t*) is the divergence-time distribution. Previously, this distribution was assumed to equal 1, but other possibilities can be explored. For example, if instead we assume that xenologous sequences are slowly *removed* from genomes due to deletions, the divergence times may be exponentially suppressed,

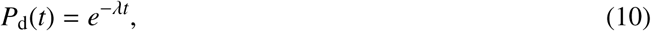

in which case equation 9 becomes:

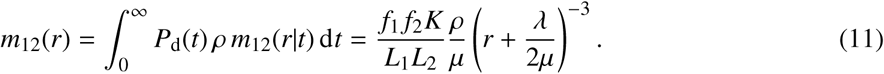

This MLD again has the familiar power-law tail in the regime *r* ≫ *λ*/(2*μ*). Generally, if the divergence-time distribution can be written as a Taylor series

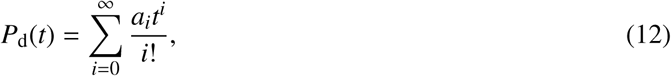

equation 9 evaluates to

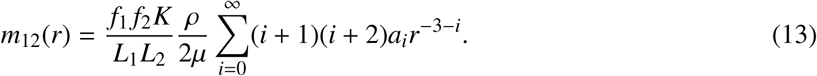

The tail of this distribution is dominated by the first nonzero term in the series, because it has the largest exponent. Again this results in a power-law with exponent −3 provided *a*_0_ = *P*_d_(0) does not vanish. That is, an exponent of −3 is expected provided HGT events have taken place at a non-zero rate up to the present time [45, 46].

### 4.5. Age-range estimation of the exact matches

According to the above model, the probability that a match of length *r* originates from an event that took place a time *t* ago is given by

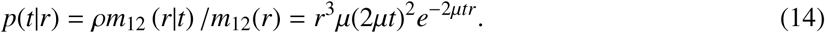

The most likely time *t*_ML_ is found by setting the time-derivative of Eq 14 to zero, which results in

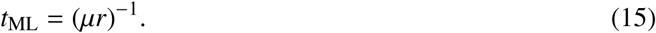

Above, we considered exact matches with a length *r* > 300 bp. Only in sequences involved in rather recent HGT events such long matches are likely to occur, and hence the method can only detect recent events. Eq 15 can provide a rough estimate for the detection horizon of the method. To do so, we substitute *r* = 300 bp into Eq 15. Assuming a mutation rate μ of 10^−9^ per bp and per generation, this results in a detection horizon of *t*_ML_ ≈ 10^6^ generations. Assuming a mean generation time in the wild of about 10 hours [28], this corresponds to approximately 1000 years. That is to say, we estimate that the HGT events we detect date back to the past 1000 years. We stress, however, that both the mutation rate and the generation time can strongly vary from one species to the next; hence this estimate is highly uncertain.

By Eq 15, the event that created the match of 19 117 bp in Fig. 1C-D is dated back about 60 years ago, again with a large uncertainty. Vancomycin was discovered in 1952, but widespread usage started only in the 1980s, and resistant strains were first reported in 1986 [39].

### 4.6. High-quality restricted dataset

To quantitatively study HGT rate variations, we restricted our analysis to a smaller and high quality dataset (see Methods) to reduce the risk of potential artefacts. The curated dataset encompasses only the exact sequence matches that stem from the comparison of contigs larger than 10^6^bp, since short contigs are more likely to present assembly or species assignment errors, or to originate from plasmid DNA. The resulting dataset comprises 138, 273 matches longer than 300bp.

We analysed exact sequence matches longer than 300bp between bacteria from different bacterial families. Here we filter out all contigs smaller than 10^6^bp from the RefSeq database. For some organisms we suspect an erroneous taxonomic annotation, due to their high similarity to another species. Based on this we manually cleaned the results and removed exact matches between following accession numbers or groups of accession numbers with particular taxonomic annotation:

- Accession number NZ_FFHQ01000001.1 and all *Enterococcus*
- Accession number NZ_JOFP01000002.1 and accession number NZ_FOTX01000001.1
- Accession number NZ_LILA01000001.1’ and all *Bacillus*
- Accession number NZ_KQ961019.1’ and all *Klebsiella*
- Accession number NZ_LMVB01000001.1’ and all *Bacillus*
- Pairwise comparisons between accession number NZ_BDAP01000001.1, NZ_JNYV01000002.1 and NZ_JOAF01000003.1

This resulted in 138, 273 unique matches.

### 4.7. Environment, Gram and GC content annotation

Ecological annotation of bacterial genera is not well defined, and different members of the same genus can occupy different ecological niches. Nevertheless, using the text mining engine of Google we annotated some of the genera as predominately Marine, Gut and Soil. Using the same approach we identified Gram-positive, Gram-negative, GC-rich and GC-poor Genera. The results are summarised in file Supp. file 7.

Additional information about bacterial genomes (such as Gram classification or lifestyle) were collected from PATRIC database metadata [75].

### 4.8. Gene enrichment analyses

To assess the enrichment of genes in the set of transferred sequences, we generated a set of control sequences as follow. For each match *i* present in *w*_*i*_ contigs, we randomly sampled without replacement a random sequences from each of those *w*_*i*_ contigs. This way, the control set takes into account the enrichment of certain species in the set of transferred sequences.

We analysed 12 different sets of genes: Acquired antibiotic resistant genes (ResFinder database [77]), Antibacterial Biocide and Metal Resistance Genes Database (BacMet database [56]), Integrative and conjugative elements(ICEberg database [3]), Virulence factors(VFDB database [11]), Essential genes (DEG database [41]), Toxin-Antitoxin systems (TADB database [68]), Peptidases (MEROPS database [64]), Bacterial Exotoxins for Human (DBETH database [10]), Transmembrane proteins (PDBTM database [36]), Restriction Enzymes (REBASE database [66]), Bacterial small regulatory RNA genes (BSRD database [40]), the Transporter Classification Database (TCDB [67]) and Enzyme classification database (Brenda [58]).

For each set of genes from a database, using the blast toolkit [1], we calculate the total number of unique match-gene hit pairs. We weighted each hit to the database by *w*_*i*_ to obtain a total number of hits *H*:

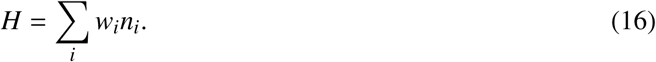

Assuming random sampling or organisms, the standard error of *H* is given by

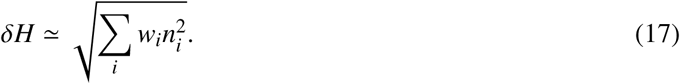

### 4.9. SEED Subsystems ontological classification

To connect identifiers of the SEED Subsystems [55] to accession identifiers of NCBI nr database, two databases were downloaded: nr from NCBI [14] FTP and m5nr from MG-RAST [47] FTP servers (on 17 January 2017). The homology search of proteins of nr database against m5nr was made using diamond [8]. Proteins from the databases were considered to have similar function if they shared 90% of amino acid similarity over the full length. Additional files for SEED Subsystems (ontology_map.gz, md5_ontology_map.gz, m5nr_v1.ontology.all) were downloaded from MG-RAST FTP.

To annotate exact matches, open reading frames were predicted with Prodigal [32] and queried against nr using diamond. After that Subsystems classification was assigned to predicted proteins when possible.

To test for enrichment we conducted the “conditional test” [59]. Briefly, it assumes that the number of “successes” in the two conditions (*H*_1_ and *H*_2_, for real data versus control) were sampled from Poisson distributions with parameters *λ*_1_ and *λ*_2_, respectively. Under the null hypothesis *λ*_1_ = *λ*_2_ the conditional probability for *H*_1_ given *H*_1_ + *H*_2_ is the binomial distribution with parameters *p* = *λ*_1_/(*λ*_1_ + *λ*_2_) = 1/2 and *n* = *H*_1_ + *H*_2_. The test is simply a binomial test that determines whether the hypothesis *p* = 1/2 can be rejected. In addition, using the Clopper–Pearson method [13], a 95% confidence interval was obtained for *p*, which was converted to a confidence interval for the enrichment *λ*_1_/*λ*_2_.

