## Supplementary Figures and Tables for "Long identical sequences found in multiple bacterial genomes reveal frequent and widespread exchange of genetic material between distant species"

### Supplementary Material

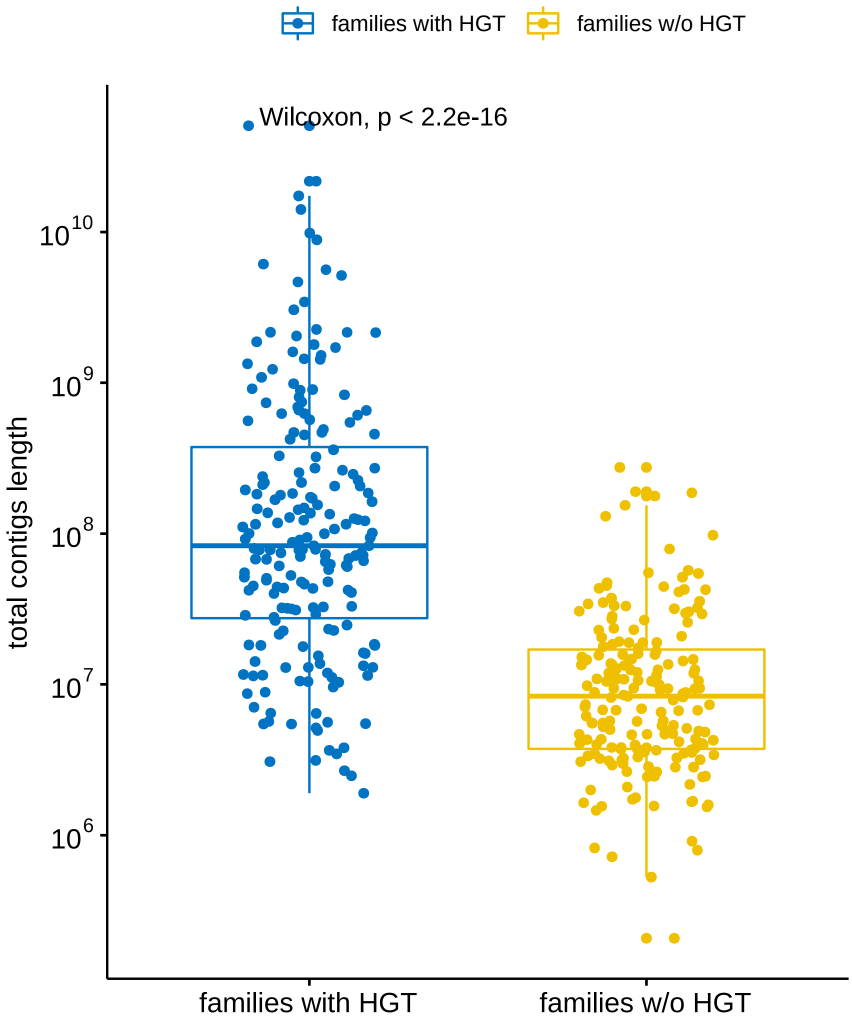


Figure S1: Total contig length distribution of families depending on their involvement in long distance HGT events. Blue: Families involved in at least one HGT event with another family. Yellow: Families involved in no long distance HGT event. The total contig length of a family is defined as the sum of the length of all the contigs belonging to that family.

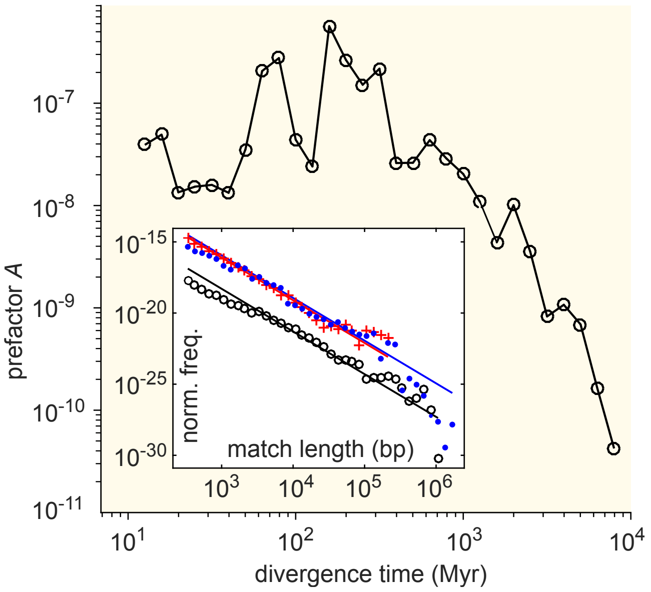


Figure S2: Effective HGT rate depending on the divergence time between genera. MLDs prefactor, $A$, resulting from comparison of genera with a given divergence time (binned) obtained from [37] in units of Myr. Inset: MLD for divergence times in the intervals $10^1 - 10^2$Myr (blue dots), $10^2 - 10^3$Myr (red pluses) and $10^3 - 10^4$Myr (black circles). Lines represent $r^{-3}$ dependence.

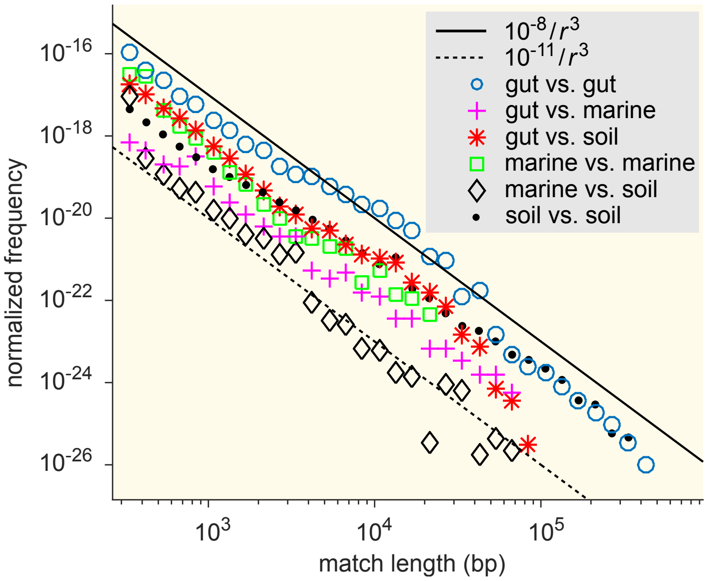


Figure S3: MLDs resulting from comparison of sets of genera associated with different ecological environments: gut, soil and marine (see Supp. file 7 for detailed annotation).

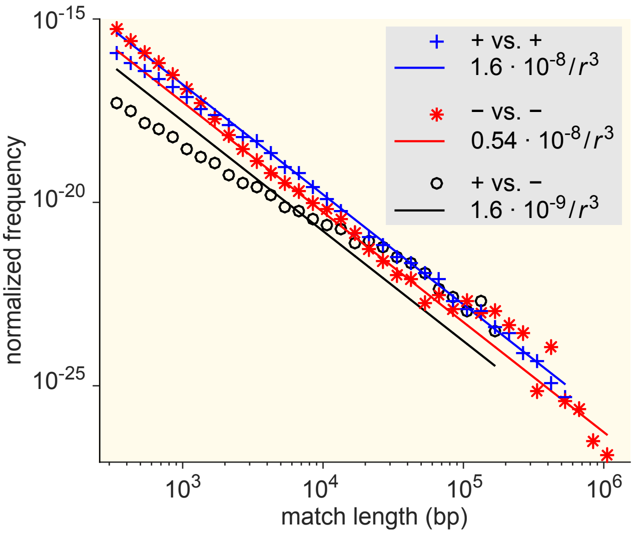


Figure S4: MLDs resulting from comparison of sets of genera associated with different gram staining test results (see Supp. file 7 for detailed annotation).

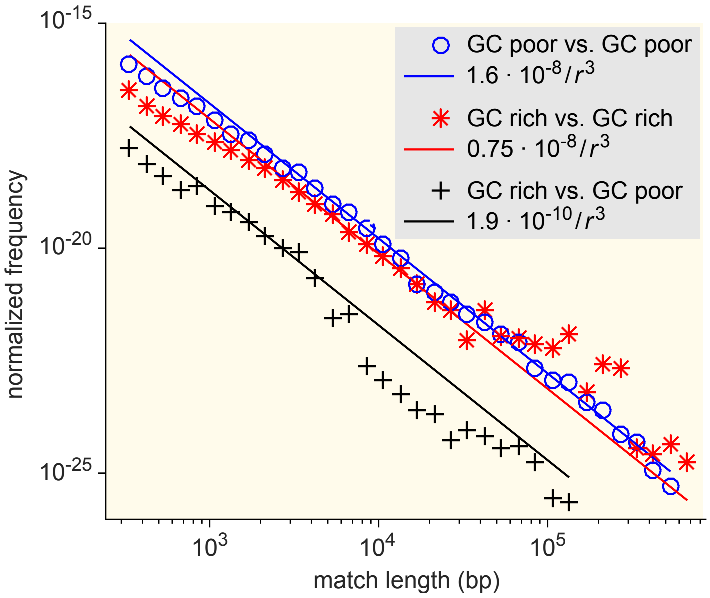


Figure S5: MLDs resulting from comparison of sets of bacteria associated with different GC content (see Supp. file 7 for detailed annotation).

| Database | Enrichment |
|---|---|
| Antimicrobial resistance [77] | 36.6 ± 0.2 |
| Biocide- and metal-resistance [56] | 3.6 ± 0.03 |
| Restriction Enzymes [66] | 0.34 ± 0.01 |
| Transmembrane Proteins [36] | |
| $\alpha$ | 0.6 ± 0.02 |
| $\beta$ | 0.98 ± 0.07 |
| Peptidases [64] | 0.084 ± 0.004 |
| ExoToxins [10] | 0.006 ± 0.003 |
| Integrative, conjugative [3] | 23.9 ± 0.1 |
| Virulence factors [11] | 1.2 ± 0.026 |
| Essential Genes [41] | 0.23 ± 0.002 |
| Small regulatory RNAs [40] | 0.01 ± 0.005 |
| Toxin-antitoxin [68] | 9.5 ± 0.6 |
| Transport Proteins [67] | |
| Channels/Pores | 0.32 ± 0.01 |
| Electrochemical Transporters | 0.56 ± 0.01 |
| Primary Active Transporters | 0.96 ± 0.01 |
| Group Translocators | 0.018 ± 0.002 |
| Transmembrane Electron Carriers | 0.026 ± 0.004 |
| Accessory Factors | 0.08 ± 0.01 |
| Incompletely Characterized | 0.24 ± 0.01 |
| Enzymes [58] | |
| Isomerases | 0.61 ± 0.001 |
| Hydrolases | 0.39 ± 0.0004 |
| Ligases | 0.35 ± 0.001 |
| Oxidoreductases | 0.33 ± 0.0004 |
| Transferases | 0.33 ± 0.0004 |
| Lyases | 0.16 ± 0.0003 |

Table S1: Enrichment of different gene categories relative to the control set (see Methods section).
